# Tgfbr2 in dental pulp cells guides sensory innervation in developing teeth

**DOI:** 10.1101/2020.04.21.053603

**Authors:** Courtney Barkley, Rosa Serra, Kevin Nguyen, Sarah B. Peters

## Abstract

Transforming growth factor β (TGFβ) plays an important role in tooth morphogenesis and mineralization. Our laboratory established a mouse model in which Tgfbr2 was conditionally deleted in the dental pulp (DP) mesenchyme using an Osterix promoter-driven Cre recombinase (Tgfbr2^cko^). These mice survived postnatally, but had significant defects in bones and teeth, including reduced mineralization and short roots. H&E staining revealed reduced axon-like structures in the mutant mice. Reporter imaging demonstrated that Osterix-Cre activity was localized within the tooth and was active in the DP and derivatives, but not in neurons. Previous research has established that paracrine signals from the dental pulp attract trigeminal axons toward and into the tooth during postnatal development, yet very little is known about the signals that regulate tooth innervation. Immunofluorescent staining for a neuronal marker, β3 tubulin, was performed on serial cryosections from control and mutant molars at two different postnatal stages. Confocal imaging and pixel quantification of β3 tubulin demonstrated reduced innervation in P7 and P24 Tgfbr2^cko^ first molars compared to controls, indicating that the signals necessary to promote neurite outgrowth were disrupted by Tgfbr2 deletion. Quantitative real time PCR showed that the mRNA expression levels of several neuronal genes were reduced. Lastly, trigeminal neurons were co-cultured atop Transwell filters overlying primary Tgfbr2^f/f^ DP cells. Tgfbr2 in the DP was deleted with an adenovirus expressing Cre recombinase. Confocal imaging of axons through the filter pores showed increased axonal sprouting from neurons cultured in the presence of Tgfbr2-positive DP cells compared to neurons cultured alone. Further, axon sprouting was reduced when Tgfbr2 was knocked down in the DP cells. These results indicate that Tgfbr2 in the DP mesenchyme regulates paracrine signals that guide sensory innervation of the tooth.

*Note: A list of abbreviations is included in the appendix*.

## Introduction

The TGFβ superfamily regulates many developmental processes, including those that occur in the tooth. Tooth development requires the coordination of epithelial, mesenchymal, hematopoietic, and neuronal cell populations to construct a functional sensory organ. During postnatal development, afferent axons from the trigeminal (TG) nerve bundle penetrate into and throughout the tooth around the time dentin deposition begins (reviewed in (Pagella et al. 2014)). This sensory innervation provides signals crucial to daily oral activities, such as eating and talking, and provides pain signals to protect the tooth organ. The roles of TGFβ signaling in specific aspects of tooth development and pathology were previously documented in mice with mutations or targeted deletions of genes encoding components of TGFβ signaling pathways (Ko et al. 2007; Zhao et al. 2008; Wang et al. 2013; Peters et al. 2017). However, Tgfbr2 deletion in the dental and skeletal mesenchyme is embryonic or perinatal lethal (Ito et al. 2003; Seo and Serra 2007; Seo and Serra 2009; Iwata et al. 2012). Since axons from the trigeminal ganglion do not penetrate the dental pulp (DP) until around postnatal day 3 (P3) (Moe et al. 2012), this lethality prevented research into the roles of TGFβ signaling in tooth innervation.

Our laboratory established an Osterix promoter-driven Cre recombinase mouse model in which *Tgfbr2* was conditionally deleted in osteoblast- and odontoblast-producing mesenchyme (*Tgfbr2*^cko^). These *Tgfbr2*^cko^ mice did not demonstrate skeletal or dental abnormalities at birth, but began to demonstrate reduced growth and mineralization in bones and teeth during the first week of life that worsened with development. The mutant phenotype included stunted root elongation and hypomineralization in molars (Wang et al. 2013; Peters et al. 2017). We stained postnatal day 7 (P7), P10, and P14 mandibular first molars (M1s) with hematoxylin and eosin (H&E) to investigate the short tooth root phenotype and noticed a reduction in axon-like structures in the mutant mice. Since Osterix-Cre should be restricted to the mesenchymal lineage (Rodda and McMahon 2006), neuronal alterations were unexpected. However, previous studies indicated that DP cells secrete neurotrophic factors to regulate tooth innervation (Kettunen 2005; Kettunen et al. 2007; Moe et al. 2012; Sijaona et al. 2012; de Almeida et al. 2014) although little information was available regarding the signaling pathways that guide this phenomenon. Furthermore, many of these studies focused on embryonic stages when the DP secretes repellants to prevent neurite extensions from entering the tooth rather than postnatal stages when the DP secretes axonal attractants (Lillesaar and Fried 2004; Fried et al. 2007). Since Osterix-Cre becomes active in the DP mesenchyme late in embryonic development (Kim et al. 2015), we hypothesized that *Tgfbr2* in the DP regulates paracrine signals guiding postnatal tooth innervation. We herein demonstrate that there is reduced innervation in *Tgfbr2*^cko^ molars compared to controls. qPCR on RNA extracted from P7 first molars indicated decreased expression of several neuronal genes. Neurite outgrowth was also reduced when *Tgfbr2* was deleted in DP cells in co-cultures of TG neurons with primary DP. Together, these data reveal a novel function for *Tgfbr2* in the development of sensory teeth.

## Materials and Methods

### Mice

All experiments with mice were approved by the UAB Institutional Animal Care and Use Committee. The Osx-Cre and *Tgfbr2*^cko^ mice were described previously (Wang et al. 2013; Peters et al. 2017). Trigeminal neuron bundles for co-cultures were collected from 6-week-old B6.Cg-Tg(Thy1-YFP)16Jrs/J mice, which express high levels of yellow fluorescent protein (YFP) in neurons (Caroni 1997). The Cre reporter strain Gt(ROSA)26SOR^tm4(ACTB-tdTomato,-EGFP)^ Luo/J (ROSA26mTmG) was crossed with Osx-Cre mice to generate Osx-Cre; ROSA26-mTmG mice with traceable Cre activity (Muzumdar et al. 2007). There were no obvious differences between males and females at P7, allowing for sample pooling for qPCR. Equivalent numbers of both sexes were used for P24 samples.

### mRNA sequence analysis

The root and crown DP from the mandibular M1s of P7 control and *Tgfbr2*^cko^ mice were collected and isolated (n=1 for each genotype). Platform information and raw data are available in the Gene Expression Omnibus (GEO; accession number GSE121285).

### Quantitative and semi-quantitative real-time PCR

The genotype for every DP sample was first confirmed. Samples from 5 controls and 5 Tgfbr2^cko^ mice were pooled for each genotype, and the experiment was repeated six times, totaling approximately 30 mice per genotype, or 60 mice total. (see Supplementary Figure 1 for a diagram showing the procedure). RNA was extracted, converted to cDNA, then the PowerUp SYBR Green Master Mix Kit (Thermo Scientific) was used for quantitative real-time polymerase chain reaction (qPCR; primers shown in Appendix Table 1). Data were normalized to Beta-2-microglobulin and analyzed with the REST software (Pfaffl et al. 2002).

**Table 1.**
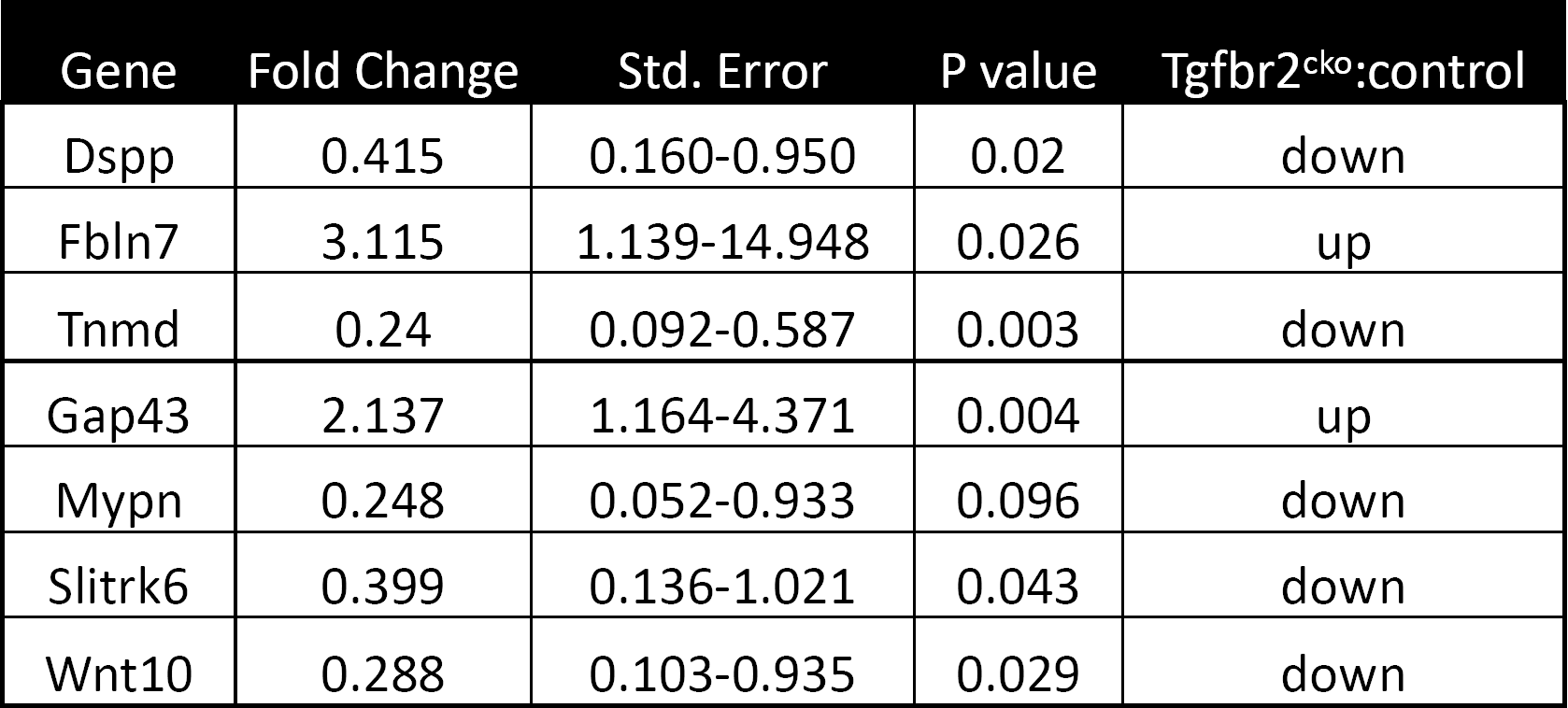
Select genes dysregulated by Tgfbr2 deletion in the P7 DP. N=6 separate analyses of 5-10 pairs of mandibular molars pooled in each sample, totaling 30 mice per genotype (control/Tgfbr2^cko^).

| Gene | Fold Change | Std. Error | P value | Tgfbr2<sup>cko</sup>:control |
| --- | --- | --- | --- | --- |
| Dspp | 0.415 | 0.160-0.950 | 0.02 | down |
| Fbln7 | 3.115 | 1.139-14.948 | 0.026 | up |
| Tnmd | 0.24 | 0.092-0.587 | 0.003 | down |
| Gap43 | 2.137 | 1.164-4.371 | 0.004 | up |
| Mypn | 0.248 | 0.052-0.933 | 0.096 | down |
| Slitrk6 | 0.399 | 0.136-1.021 | 0.043 | down |
| Wnt10 | 0.288 | 0.103-0.935 | 0.029 | down |

*Tgfbr2* knockdown in the co-culture model was confirmed using our previously-developed semi-qPCR assay (Seo and Serra 2007), and the results were confirmed in three separate experiments.

### Imaging of DP innervation

Mandibles were fixed in paraformaldehyde for one hour, followed by decalcification. Tissue was embedded, frozen and sectioned on the sagittal plane at 10 μm thickness with a cryostat. For immunofluorescence, sections were permeabilized with cold acetone at -20°C for 10 minutes and rinsed twice with phosphate-buffered solution (PBS). Slides were immersed in hot citrate buffer for antigen retrieval then rinsed in PBS. Histology slides were similarly immersed in citrate buffer and stained with hematoxylin and eosin. Immunofluorescence slides were blocked with 5% bovine serum albumin (BSA), 5% goat serum and Mouse-on-Mouse blocking serum (Vector Labs) in PBS for 1 hour at room temperature, followed by overnight incubation at 4°C in anti-β3 tubulin (1:100, R&D Systems). Sections were rinsed and incubated with goat-anti-mouse Alexa-488 (1:500, Invitrogen) and DAPI (1:1000, ThermoScientific) overnight at 4°C. Ten-micron thick z-stacks were collected and presented in maximum projection mode, quantified for total pixels and normalized to the DAPI pixels using FIJI software. Unpaired t-tests were performed using samples from four separate mice of each genotype (P < 0.05 considered significant).

### Trigeminal neuron and DP primary co-culture

Co-culture assays were performed as described previously (Barkley et al. 2020). Briefly, primary DP cells from P5-8 *Tgfbr2*^fl/fl^ molars were grown in alpha-minimum essential medium (alpha-MEM; Gibco) supplemented with 10% heat-inactivated fetal bovine serum, L-glutamine (Gibco), and antibiotics. At confluence, the media were replaced with media containing Adenovirus-Cre-GFP or Adenovirus-eGFP for 48 hours before replacement with fresh media. Cells from dispersed TGN bundles (50,000) from 6-week-old B6.Cg-Tg(Thy1-YFP)16Jrs/J male and female mice were plated on Transwell inserts (Grenier Thincerts, 3 μm porosity) coated with 10 μg/ml laminin (Sigma). Mitotic inhibitors, 1 mM uridine (Sigma) and 15 mM 5’-Fluor-2’deoxyuridine (Sigma), were added after 24 hours. Co-culture continued for an additional 4 days, then filters containing TGN were fixed, blocked and incubated with anti-GFP primary antibody (1:200, Aves Lab) to selectively stain the YFP+ neurons (due to the high conservation between GFP and YFP). Filters were subsequently rinsed and incubated in Alexa-546 secondary antibody (1:500, Invitrogen). Finally, 100 μm thick z-stacks were stitched together in NIS-Elements and presented in maximum projection mode to demonstrate total neurite outgrowth throughout the entire filter. Images were thresholded with FIJI software with the Intermodes autothreshold function and pixels were quantified. Two-tailed ratio-paired t-tests were performed for four separate co-culture experiments (significance set to P<0.05).

## Results

### Tgfbr2 deletion is isolated to the DP mesenchyme

*Tgfbr2*^cko^ mice demonstrated alterations in development of the mandibular M1s by P5 (Wang et al. 2013). To investigate the mechanisms responsible for the phenotypes observed in *Tgfbr2*^cko^ teeth, we performed H&E staining on sections from P7, P10 and P14 control and mutant M1s. These time points were chosen because tooth root elongation and dentin mineralization were both actively occurring. Axon-like structures were reduced in mutant teeth relative to controls at all stages (Supplementary Figure 1). We assessed whether the hypothesized reduction of innervation in the Tgfbr2^cko^ mice was due to unexpected Cre activity in neurons by crossing Osterix-Cre and ROSA26-mTmG reporter mice. The ROSA26-mT/mG reporter line expresses a membrane-bound GFP (mG) tag in cells that express Cre at any point, or a Tomato red fluorescent (mT) tag in cells that never express Cre (Muzumdar et al. 2007). Osterix:GFP::Cre is also expressed in the nucleus (Rodda and McMahon 2006). We collected P7 Osterix-Cre;ROSA26mTmG DP and performed immunofluorescence staining for β3T on whole mount DP with a Cy5 secondary antibody, followed by confocal microscopy for mG, mT, and Cy5 to detect neurons (Figure 1). The DP mesenchyme expressed GFP, indicating Cre activity (Figure 1A). Yellow inset boxes indicate several mT-positive structures (Figure 1B, pseudocolored blue) that are also β3T-positive (Figure 1C, indicating axons. The asterisk indicates a larger, β3T-negative and mT-positive structure, likely a large blood vessel (Figure 1 B,C) indicating the blood vessels also did not express Cre. The β3T stained tissue did not directly co-localize with GFP (Figure 1D), indicating that Cre was not active in the axons. These results suggest that there was no off-target deletion of *Tgfbr2*.

**Figure 1.**
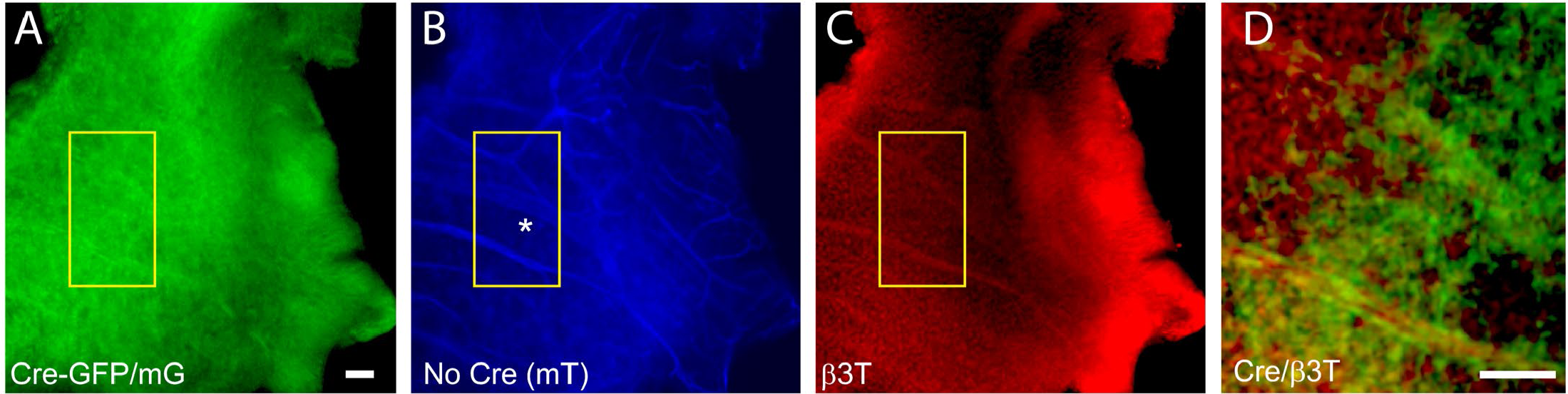
Cre activity in whole-mounted P7 Cre+;ROSA26mTmG mouse dental pulp. Whole mount dental pulp images show apical/root structures to the left of the image and crown to the right. (A) Cre expression was detected as green fluorescence due to the expression of Osterix-Cre-GFP and mG. (B) Cells in which Cre was never active are shown in mT images pseudocolored blue. (C) β3T+ staining (red) of neuronal structures localized in some areas of mT structures shown in (B). The asterisk in (B) indicates an mT structure that does not stain for β3T in (C), labelling a hypothesized blood vessel. The zoomed-in area in (D) is the area indicated with the yellow box in A-C. The merged staining shows that the axonal structures were distinctly red, without the green co-localization of Cre+ mesenchyme. N=7. The scale bars are 50 μm for A-C and 50 μm for D.

### Afferent innervation is reduced in Tgfbr2^cko^ molars

To directly assess innervation, sections from P7 and P24 control and mutant molars were stained with pan-neuronal β3T, and the levels of fluorescence were compared. Nerve fascicles branched throughout both the control and mutant DP, with individual axons entering the predentine and dentine regions of the tooth organ (Figure 2). Maximum projections from confocal microscopy were used to evaluate and quantify fluorescence as a measure of tooth innervation. Total axon coverage was quantified as the pixel density of equivalent regions of the image normalized to DAPI. We found a significant reduction in nerve fiber coverage in molars from P7 and P24 *Tgfbr2*^cko^ mice compared to controls (Figure 2) supporting the hypothesis that *Tgfbr2* in the DP mesenchyme mediates tooth innervation. Since *Tgfbr2*^cko^ mice die around 4 weeks of age (Wang et al. 2013; Peters et al. 2017), we were unable to investigate beyond this timepoint.

**Figure 2.**
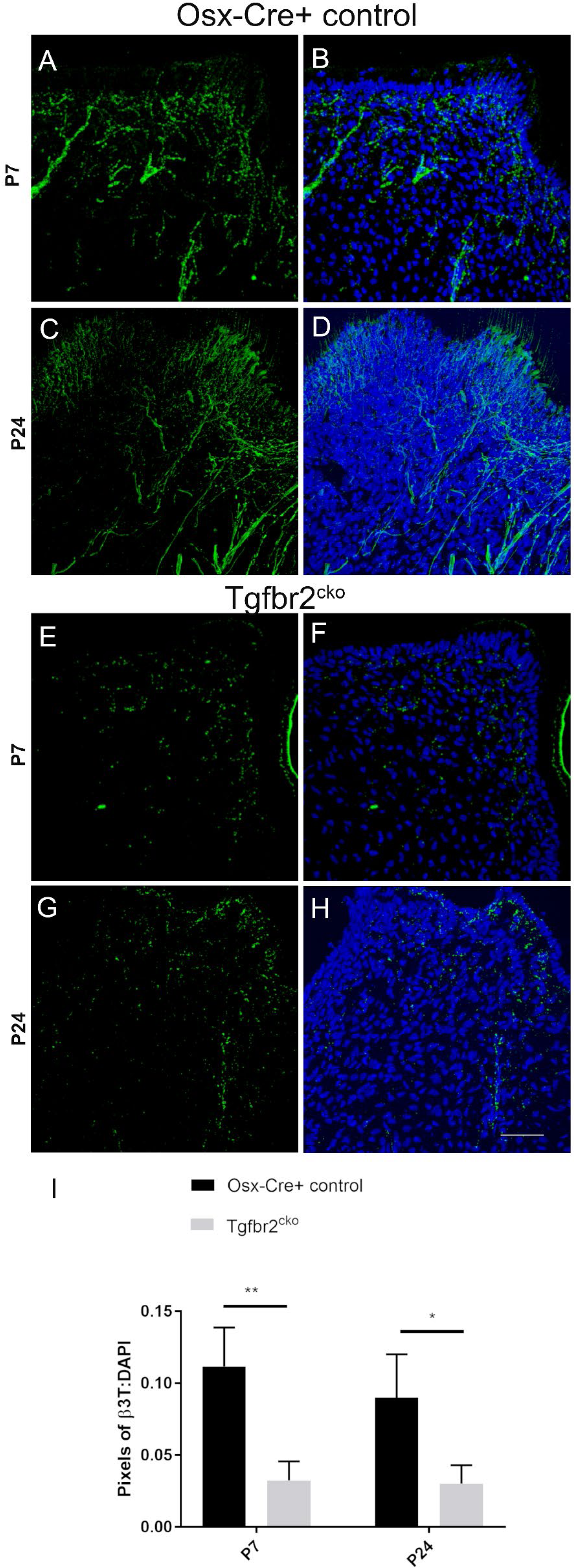
DP innervation in control and *Tgfbr2*^cko^ P7 and P24 first molars. Z-stacks (10 μm thick) were collected and presented in maximum projection mode to examine the total afferent innervation. Confocal images of control (A-D) and *Tgfbr2*^cko^ (E-H) DP demonstrated more β3T+ neuronal structures in control mice than mutant mice. The scale bar is 50 μm. (E) The ratios of total pixels for neuronal structures normalized to the pixels for DAPI staining indicate significantly more innervation per region in control mice than mutant mice. *, p< 0.05, ** p<0.01 by unpaired t-tests, n=4 for each genotype at each time point.

### Expression of select genes is altered in Tgfbr2^cko^ molars

We next assessed the expression of several genes, including neuronal genes, identified by a preliminary mRNA Sequence Analysis on mRNA from P7 root and crown DP samples (GEO; accession number GSE121285) by performing qPCR on P7 DP from control and mutant mice, shown in Table 1. We verified the mutant phenotype by demonstrating decreased expression of *Dspp* and *Tnmd* in the mutant samples, reflecting the hypomineralization and stunted root growth, respectively (Pryce et al. 2009; Wang et al. 2013). There was also increased expression of Fibulin-7 (*Fbln7*), a transcript encoding for a cellular adhesion proteins in the dentin matrix (Sarangi et al. 2018). For neuronal transcripts, qPCR demonstrated upregulation of growth-associated protein 43 (*Gap43*) (Curtis et al. 1992; Jessen and Mirsky 2005), which was verified by immunostaining and confocal microscopy (Supplemental Figure 3). There was also downregulation of neuronal genes SLIT and NTRK Like Family Member 6 (*Slitrk6*) (Katayama et al. 2009), Wnt Family Member 10a (*Wnt10a*) (Matsukawa et al. 2018) and myopalladin (*Mypn*) (Otey et al. 2005), with no impact on *Gphn* (data not shown). These findings confirmed that the neuronal transcriptome was altered in mutant mouse molars.

### Neurite outgrowth in DP-TGN co-cultures requiresTgfbr2

We utilized co-cultures to directly assess the role of Tgfbr2 in neurite outgrowth in response to DP secretions. Transwell filters were seeded with TGN while DP cells were grown in the bottom well. TGN from Thy1-YFP mice were used so that neurite outgrowth through the filter could be easily imaged. We observed the brightest staining of Thy1-YFP axons with the GFP antibody at 6 weeks of age, so TGN from 6-week-old mice were used in co-cultures. When the TGN were grown on filters in the absence of DP, very little neurite outgrowth occurred (Figure 4C,F). Significantly higher neurite outgrowth was observed in the presence of DP cells (Figure 4A,F). To examine whether *Tgfbr2* in the DP regulates neurite outgrowth, *Tgfbr2* was deleted using Adenovirus-Cre-GFP. Cells infected with Adenovirus-eGFP were used as controls. Equivalent numbers of infected cells were used (as determined by GFP fluorescence; Figure 4D), and the deletion of Tgfbr2 was determined by semi-quantitative PCR (Figure 4E). Deletion of *Tgfbr2* in the DP significantly decreased the neurite extension, resulting in levels similar to cultures without DP (Figure 4A-C,F). These results suggest that *Tgfbr2* in the DP mesenchyme is required to promote neurite outgrowth.

**Figure 3.**
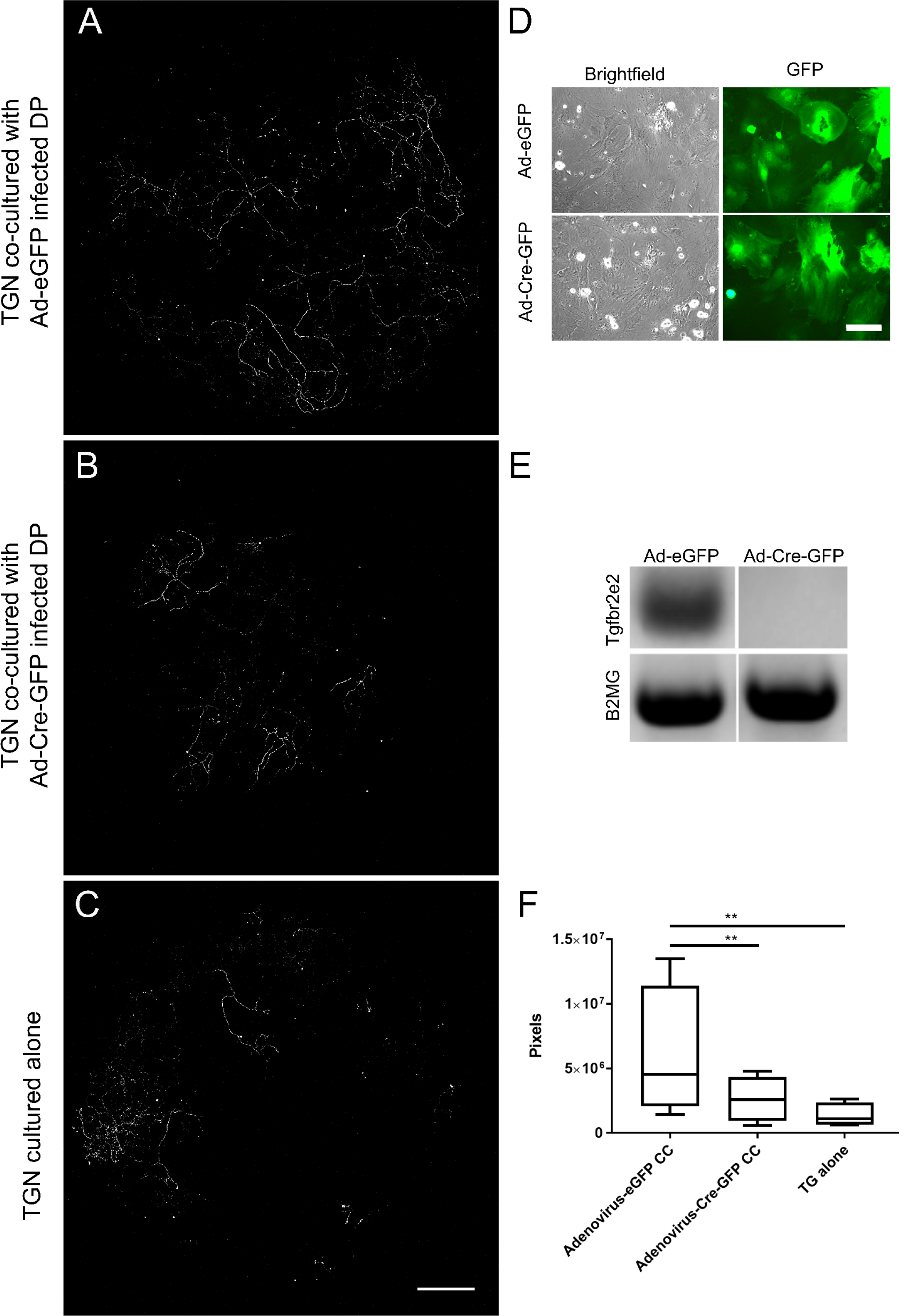
Primary DP cell signaling mediates trigeminal neurite outgrowth, apparently via Tgfbr2. (A-C) Thy1-YFP trigeminal neurons were cultured in Transwell filters with 3 μm pores atop primary *Tgfbr2*^f/f^ DP cells. Immunofluorescence was performed for the YFP protein using an anti-GFP antibody for highly specific staining of neuronal structures over the entire filter. Maximum projections of 100 μm z-stack confocal microscopy images at 10x were collected and stitched with stitching software. TG neurons demonstrated significantly more outgrowth when co-cultured with DP cells (A) than when cultured alone (C). Neurite outgrowth was not induced when neurons were co-cultured with DP infected with Adenovirus-Cre-GFP to knock down *Tgfbr2* (B, F). The scale bar is 1000 μm. Equivalent numbers of cells infected with Adenovirus-eGFP and Adenovirus-Cre-GFP are demonstrated in (D). The scale bar is 125 μm. Semi-quantitative PCR confirmed *Tgfbr2* KD (E). Images of axonal outgrowth on the filters were auto-thresholded with FIJI software to remove background fluorescence and the remaining pixels were quantified (F). n=4 separate experiments, ** P< 0.01 by two-tailed ratio-paired t-tests.

**Figure 4.**
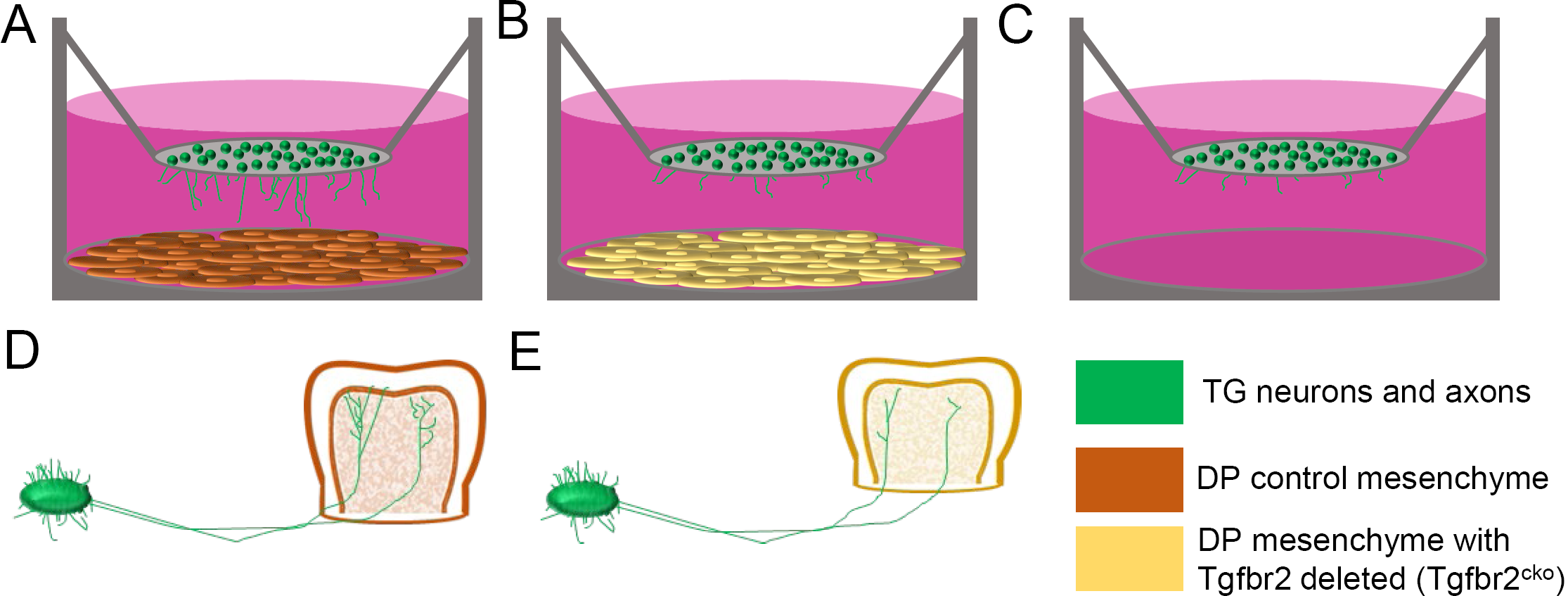
Schematic model demonstrating neurite outgrowth in response to Tgfbr2 signaling from the dental pulp mesenchyme. (A) The co-culture model of TG neurons (green) grown atop a Transwell filter overlying control DP cells (dark orange). (B) Co-culture assays were performing using TG neurons overlying DP cells with Tgfbr2 deleted (yellow). (C) TG neurons grown alone atop a Transwell filter. (D) A model of TG ganglia extending axons into and throughout a developing control molar *in vivo*. (E) A model of TG ganglia extending axons into a developing Tgfbr2^cko^ molar, exhibiting less neurite outgrowth throughout the pulp. The images are for illustration purposes only and are not drawn to scale.

In summary, deletion of *Tgfbr2* in the DP during TGN penetration and sprouting in the tooth leads to altered expression of neuronal genes and reduced innervation in postnatal teeth. Co-culture experiments showed that *Tgfbr2* in the DP is required for DP-mediated neurite outgrowth. Figure 4 recaps these data. We propose that *Tgfbr2* regulates paracrine signaling in the DP, which acts on TGN to regulate innervation of the postnatal tooth.

## Discussion

Tooth innervation serves unique roles that allow teeth to sense pressure, temperature, and infectious agents. Without these sensations, decay and normal oral activities would cause irreparable damage to the tooth and surrounding tissues. Human teeth are primarily innervated by nociceptors extending from the trigeminal ganglia (Fried et al. 2007) that respond to signals from the dental mesenchyme during development (Lillesaar et al. 1999; Fried et al. 2005; Kettunen 2005; Kettunen et al. 2007). While neuronal-mesenchymal crosstalk is required for tooth organogenesis and maintenance, the details of these mechanisms have been largely unclear. Our observation of reduced innervation in *Tgfbr2*^cko^ mice led us to investigate its role in this cross-talk. The *in vitro* co-culture assay used media lacking factors to promote either mineralization in the DP cells or growth factors to promote neurite outgrowth in order to directly correlate paracrine signals from the DP cells to neurite outgrowth in the overlying neurons. The results (summarized in Figure 4) suggest a novel role for *Tgfbr2* in the DP mesenchyme in the sensory innervation of teeth.

To elucidate the mechanisms underlying the phenotypes observed in *Tgfbr2*^cko^ teeth, we performed qPCR on P7 DP on several genes. We found reduced expression of dentin sialophosphoprotein (*Dspp*) in mutant DP, as previously noted in P10 and P14 mutant molars (Wang et al. 2013). The expression of tenomodulin (*Tnmd*), which is regulated by TGFβ signaling (Pryce et al. 2009) and expressed in the periodontal ligament covering tooth roots (Shukunami et al. 2016), was also reduced coincident with the reduced roots in the mutant mice. Of note, there was also upregulation of Fibulin-7 (*Fbln7*), a gene involved in cellular adhesion during predentin and dentin formation (de Vega et al. 2007), in the Tgfbr2^cko^ dental pulp compared to control teeth. The increased *Fbln7* in the Tgfbr2^cko^ dental pulp may be present to compensate for the decreased odontoblast differentiation and dentin mineralization in these mice (Wang et al. 2013). Additional studies are needed to elucidate how *Fbln7* contributes to the cell-extracellular matrix interactions during tooth development.

We then performed qPCR for neuronal genes where down-regulation was likely a consequence of reduced innervation. For instance, *Mypn* was recently identified as a marker of axon terminals (Woods et al. 2018). We found that *Mypn* trended toward downregulation, which corresponded to the reduction of afferents in β3T images (Figure 2). Neurite outgrowth and axon elongation occur through the growth cones at the distal tips of growing axons. Growth-associated protein 43 (Gap43) is expressed in growth cones during development (Meiri et al. 1986; Goslin et al. 1988). However, Gap43-deficient mice still have axonal outgrowth, indicating that Gap43 is not required for axonal growth. Still, Gap43-deficient axons were unable to navigate past certain decision points, suggesting that Gap43 may provide pathfinding signals to the growth cones. Interestingly, *Gap43* is also widely expressed in Schwann cell precursors (Jessen and Mirsky 2005) and mature Schwann cells (Curtis et al. 1992) of developing nerves, and its expression may increase with reduced innervation (Curtis et al. 1992). Our immunofluorescent images of Gap43 (Supplemental Figure 3) showed a distinct morphology in Gap43-containing structures in the Tgfbr2^cko^ molars that resemble Schwann cells, whereas the structures in the control mice were more like axonal afferents. We hypothesize that the increase in *Gap43* expression observed in our samples may have been from Schwann cells responding to the lack of axonal contact. Further studies will be needed to clarify how the DP mesenchyme affects axon pathfinding through Gap43 expression in either/both axons and Schwann cells.

Previous studies demonstrate that the DP mesenchyme secretes neurotrophic factors, which we further investigated with our qPCR analyses. Slitrk6 is expressed in the developing tooth mesenchyme (Aruga 2003) and is known to promote the innervation of organs by regulating non-neuronal expression of neurotrophins (Katayama et al. 2009). Our qPCR data indicate that there was decreased expression of Slitrk6 in the mutant mice, suggesting that a feedback mechanism, rather than direct suppression of neurotrophin secretion, could be disrupted. qPCR also supported *Tgfbr2*-mediated regulation of *Wnt10a*, which encodes a secreted signaling protein, in the DP. *Wnt10a* loss is associated with tooth agenesis and abnormal tooth formation (OMIM #257980, 224750, 150400). A recent study evaluated the role of Wnt ligands in odontoblast differentiation and dentin formation by disrupting the chaperon protein, Wntless, which regulates Wnt sorting and secretion. The mutant mice demonstrated a phenotype similar to *Tgfbr2*^cko^ mice, with decreased dentin thickness and shortened tooth roots (Bae et al., 2015). Wnt proteins are also known to play important roles in the nervous system, and Wnt10a is thought to promote axon growth by activating small G-proteins (Matsukawa et al., 2018). In addition, the mature distribution of axons does not occur until root formation is nearly complete (Fristad et al., 1994), and this did not occur in the mutant mice. Future studies could create and utilize a Wnt10^f/f^ mouse model to conditionally delete this gene in mesenchymal or neuronal progenitors to fully investigate the roles of each population in tooth root development. We believe these studies would be of particular interest to the field of orthodontics.

Our qPCR and co-culture results suggest that axonal growth occurs in response to a combination of paracrine signals, with Tgfbr2 being required for neurite outgrowth. Further research is necessary to determine which signals and the precise timing required to promote axonogenesis and/or guide the axons to their appropriate targets during tooth innervation. A previous study demonstrated that the conditional deletion of Tgfbr2 in odontoblasts led to disrupted matrix secretion that calcified and obliterated the pulp chamber with age (Ahn et al. 2015). This indicated that Tgfbr2 is crucial to maintaining soft pulp tissue, including the neuronal structures contained in it. In mature teeth, axons sprout after tooth injury and then signal to the DP to repair the damaged dentin (Taylor et al. 1988). We suggest that there is a neuronal-mesenchymal feedback loop guiding tooth maintenance and that activation of TGFβ pathways could stimulate axonal sprouting in regions of tooth damage. Future studies will utilize the inducible (tet-off) Osterix-GFP::Cre system (Wang et al. 2013) to study whether TGFβ-guided signals in the mature DP are necessary and sufficient to repair tooth injuries.

Finally, while the results of this study have clear implications for the fields of pediatric dentistry and regenerative endodontics, we believe that the present findings might also have applications to other fields, such as stroke and spinal cord or nerve damage.

## Figure and Table Legends

**Supplemental Figure 1. Supplemental Figure 1**. Schematic of DP RNA collection and processing for qPCR analysis. (A) 5 samples of control are pooled. (B) 5 samples of Tgfbr2^cko^ samples are pooled. (A+B) are considered an n of 1. (C) Chart demonstrating 12 independently pooled samples for 6 separated analyses. Each mouse/sample is assigned a number. (D) qPCR is performed for 6 biological replicates for each genotype.

**Supplemental Figure 2**. Nerve structures in postnatal teeth are reduced in Tgfbr2^cko^ mice. Representative images of H&E-stained mandibular M1s from postnatal days 7, 10, and 14. The black arrows in each image indicate hypothesized neuronal structures. Several other structures are present, but are not indicated to prevent image crowding. N=3 for each genotype. Scale bar = 100 μm.

**Supplemental Figure 3**. Gap43 expression is increased in Tgfbr2^cko^ mice. Z-stacks (10 μm thick) were collected and presented in maximum projection mode to examine Gap43 expression. Representative confocal images of control (A, B) and Tgfbr2^cko^ (C, D) dental pulp demonstrated more Gap43 in the mutant mice. The scale bar is 25 μm. n=5 per genotype.

## Acknowledgements

This work was supported by the a) National Institutes of Health/NIAMS grant numbers R01 AR062507 and R01 AR053860 to RS, b) the University of Alabama at Birmingham Dental Academic Research Training (DART) grant T90DE022736 (PI MacDougall) to SBP from the National Institute of Dental and Craniofacial Research/National Institutes of Health, c) UAB Global Center for Craniofacial, Oral and Dental Disorders (GC-CODED) Pilot and Feasibility grant to SBP and d) the National Institute of Dental and Craniofacial Research/National Institutes of Health K99 DE024406 grant to SBP.

We would like to thank Drs. Michael Crowley and David Crossman from the UAB Heflin Center for Genomic Sciences for their assistance with the mRNA Sequence Analysis and bioinformatics, respectively.

There are no conflicts of interest.

## References

Ahn YH, Kim TH, Choi H, Bae CH, Yang YM, Baek JA, Lee JC, Cho ES. 2015. Disruption of Tgfbr2 in odontoblasts leads to aberrant pulp calcification. J Dent Res. 94(6):828–35. doi:10.1177/0022034515577427. http://www.ncbi.nlm.nih.gov/pubmed/25818583.

de Almeida JFA, Chen P, Henry MA, Diogenes A. 2014. Stem cells of the apical papilla regulate trigeminal neurite outgrowth and targeting through a BDNF-dependent mechanism. Tissue Eng Part A. 20(23–24):3089–100. doi:10.1089/ten.TEA.2013.0347. http://www.ncbi.nlm.nih.gov/pubmed/24837134.

Aruga J. 2003. Slitrk6 expression profile in the mouse embryo and its relationship to that of Nlrr3. Gene Expr Patterns. 3(6):727–733. doi:10.1016/S1567-133X(03)00141-8. https://www.sciencedirect.com/science/article/pii/S1567133X03001418?via%3Dihub.

Bae CH, Kim TH, Ko SO, Lee JC, Yang X, Cho ES. 2015. Wntless Regulates Dentin Apposition and Root Elongation in the Mandibular Molar. J Dent Res. 94(3):439–45. doi:10.1177/0022034514567198. http://www.ncbi.nlm.nih.gov/pubmed/25595365.

Barkley C, Serra R, Peters SB. 2020. A Co-Culture Method to Study Neurite Outgrowth in Response to Dental Pulp Paracrine Signals. J Vis Exp. (156). doi:10.3791/60809. http://www.ncbi.nlm.nih.gov/pubmed/32116290.

Caroni P. 1997. Overexpression of growth-associated proteins in the neurons of adult transgenic mice. J Neurosci Methods. 71(1):3–9. doi:10.1016/S0165-0270(96)00121-5. https://www.sciencedirect.com/science/article/pii/S0165027096001215?via%3Dihub.

Curtis R, Stewart HJ, Hall SM, Wilkin GP, Mirsky R, Jessen KR. 1992. GAP-43 is expressed by nonmyelin-forming Schwann cells of the peripheral nervous system. J Cell Biol. 116(6):1455–64. doi:10.1083/JCB.116.6.1455. http://www.ncbi.nlm.nih.gov/pubmed/1531832.

Fried K, Lillesaar C, Sime W, Kaukua N, Patarroyo M. 2007. Target finding of pain nerve fibers: Neural growth mechanisms in the tooth pulp. Physiol Behav. 92(1–2):40–45. doi:10.1016/J.PHYSBEH.2007.05.032. https://www.sciencedirect.com/science/article/pii/S0031938407002272?via%3Dihub.

Fried K, Sime W, Lillesaar C, Virtanen I, Tryggvasson K, Patarroyo M. 2005. Laminins 2 (α2β1γ1, Lm-211) and 8 (α4β1γ1, Lm-411) are synthesized and secreted by tooth pulp fibroblasts and differentially promote neurite outgrowth from trigeminal ganglion sensory neurons. Exp Cell Res. 307(2):329–341. doi:10.1016/j.yexcr.2005.04.009. http://www.ncbi.nlm.nih.gov/pubmed/15894315.

Goslin K, Schreyer DJ, Skene JHP, Banker G. 1988. Development of neuronal polarity: GAP-43 distinguishes axonal from dendritic growth cones. Nature. 336(6200):672–674. doi:10.1038/336672a0. http://www.nature.com/articles/336672a0.

Ito Y, Yeo JY, Chytil A, Han J, Bringas P, Nakajima A, Shuler CF, Moses HL, Chai Y. 2003. Conditional inactivation of Tgfbr2 in cranial neural crest causes cleft palate and calvaria defects. Development. 130(21):5269–80. doi:10.1242/dev.00708. http://www.ncbi.nlm.nih.gov/pubmed/12975342.

Iwata J, Hacia JG, Suzuki A, Sanchez-Lara PA, Urata M, Chai Y. 2012. Modulation of noncanonical TGF-β signaling prevents cleft palate in Tgfbr2 mutant mice. J Clin Invest. 122(3):873–85. doi:10.1172/JCI61498. http://www.jci.org/articles/view/61498.

Jessen KR, Mirsky R. 2005. The origin and development of glial cells in peripheral nerves. Nat Rev Neurosci. 6(9):671–682. doi:10.1038/nrn1746.

Katayama K, Zine A, Ota M, Matsumoto Y, Inoue T, Fritzsch B, Aruga J. 2009. Disorganized Innervation and Neuronal Loss in the Inner Ear of Slitrk6-Deficient Mice. Sham MH, editor. PLoS One. 4(11):e7786. doi:10.1371/journal.pone.0007786. http://www.ncbi.nlm.nih.gov/pubmed/19936227.

Kettunen P. 2005. Coordination of trigeminal axon navigation and patterning with tooth organ formation: epithelial-mesenchymal interactions, and epithelial Wnt4 and Tgfb1 regulate semaphorin 3a expression in the dental mesenchyme. Development. 132(2):323–334. doi:10.1242/dev.01541. http://dev.biologists.org/cgi/doi/10.1242/dev.01541.

Kettunen P, Spencer-Dene B, Furmanek T, Kvinnsland IH, Dickson C, Thesleff I, Luukko K. 2007. Fgfr2b mediated epithelial-mesenchymal interactions coordinate tooth morphogenesis and dental trigeminal axon patterning. Mech Dev. 124(11–12):868–83. doi:10.1016/j.mod.2007.09.003. http://linkinghub.elsevier.com/retrieve/pii/S0925477307001487.

Kim TH, Bae CH, Lee JC, Kim JE, Yang X, de Crombrugghe B, Cho ES. 2015. Osterix Regulates Tooth Root Formation in a Site-specific Manner. J Dent Res. 94(3):430–438. doi:10.1177/0022034514565647. http://www.ncbi.nlm.nih.gov/pubmed/25568170.

Ko SO, Chung IH, Xu X, Oka S, Zhao H, Cho ES, Deng C, Chai Y. 2007. Smad4 is required to regulate the fate of cranial neural crest cells. Dev Biol. 312(1):435–47. doi:10.1016/j.ydbio.2007.09.050. http://www.pubmedcentral.nih.gov/articlerender.fcgi?artid=2704603&tool=pmcentrez&rendertype=abstract.

Lillesaar C, Eriksson C, Johansson CS, Fried K, Hildebrand C. 1999. Tooth pulp tissue promotes neurite outgrowth from rat trigeminal ganglia in vitro. J Neurocytol. 28(8):663–70. http://www.ncbi.nlm.nih.gov/pubmed/10851345.

Lillesaar C, Fried K. 2004. Neurites from trigeminal ganglion explants grown in vitro are repelled or attracted by tooth-related tissues depending on developmental stage. Neuroscience. 125(1):149–161. doi:10.1016/j.neuroscience.2004.01.008. http://www.ncbi.nlm.nih.gov/pubmed/15051154.

Matsukawa T, Morita K, Omizu S, Kato S, Koriyama Y. 2018. Mechanisms of RhoA inactivation and CDC42 and Rac1 activation during zebrafish optic nerve regeneration. Neurochem Int. 112:71–80. doi:10.1016/j.neuint.2017.11.004. https://linkinghub.elsevier.com/retrieve/pii/S0197018617303285.

Meiri KF, Pfenninger KH, Willard MB. 1986. Growth-associated protein, GAP-43, a polypeptide that is induced when neurons extend axons, is a component of growth cones and corresponds to pp46, a major polypeptide of a subcellular fraction enriched in growth cones. Proc Natl Acad Sci U S A. 83(10):3537–41. doi:10.1073/pnas.83.10.3537. http://www.ncbi.nlm.nih.gov/pubmed/3517863.

Moe K, Sijaona A, Shrestha A, Kettunen P, Taniguchi M, Luukko K. 2012. Semaphorin 3A controls timing and patterning of the dental pulp innervation. Differentiation. 84(5):371–379. doi:10.1016/j.diff.2012.09.003. http://www.sciencedirect.com/science/article/pii/S0301468112001314.

Muzumdar MD, Tasic B, Miyamichi K, Li L, Luo L. 2007. A global double-fluorescent Cre reporter mouse. genesis. 45(9):593–605. doi:10.1002/dvg.20335. http://doi.wiley.com/10.1002/dvg.20335.

Otey CA, Rachlin A, Moza M, Arneman D, Carpen O. 2005. The palladin/myotilin/myopalladin family of actin-associated scaffolds. Int Rev Cytol. 246:31–58. doi:10.1016/S0074-7696(05)46002-7.

Pagella P, Jiménez-Rojo L, Mitsiadis TA. 2014. Roles of innervation in developing and regenerating orofacial tissues. Cell Mol Life Sci. 71(12):2241–2251. doi:10.1007/s00018-013-1549-0. http://www.ncbi.nlm.nih.gov/pubmed/24395053.

Peters SB, Wang Y, Serra R. 2017. Tgfbr2 is required in osterix expressing cells for postnatal skeletal development. Bone. 97:54–64. doi:10.1016/j.bone.2016.12.017.14]. http://linkinghub.elsevier.com/retrieve/pii/S8756328216303878.

Pfaffl MW, Horgan GW, Dempfle L. 2002. Relative expression software tool (REST) for group-wise comparison and statistical analysis of relative expression results in real-time PCR. Nucleic Acids Res. 30(9):e36. http://www.pubmedcentral.nih.gov/articlerender.fcgi?artid=113859&tool=pmcentrez&rendertype=abstract.

Pryce BA, Watson SS, Murchison ND, Staverosky JA, Dünker N, Schweitzer R. 2009. Recruitment and maintenance of tendon progenitors by TGFbeta signaling are essential for tendon formation. Development. 136(8):1351–61. doi:10.1242/dev.027342. http://www.ncbi.nlm.nih.gov/pubmed/19304887.

Rodda SJ, McMahon AP. 2006. Distinct roles for Hedgehog and canonical Wnt signaling in specification, differentiation and maintenance of osteoblast progenitors. Development. 133(16):3231–44. doi:10.1242/dev.02480. http://dev.biologists.org/content/133/16/3231.

Sarangi PP, Chakraborty P, Dash SP, Ikeuchi T, de Vega S, Ambatipudi K, Wahl L, Yamada Y. 2018. Cell adhesion protein fibulin-7 and its C-terminal fragment negatively regulate monocyte and macrophage migration and functions *in vitro* and *in vivo*. FASEB J. 32(9):4889–4898. doi:10.1096/fj.201700686RRR. https://www.fasebj.org/doi/10.1096/fj.201700686RRR.

Seo H-S, Serra R. 2007. Deletion of Tgfbr2 in Prx1-cre expressing mesenchyme results in defects in development of the long bones and joints. Dev Biol. 310(2):304–16. doi:10.1016/j.ydbio.2007.07.040. http://www.pubmedcentral.nih.gov/articlerender.fcgi?artid=2042108&tool=pmcentrez&rendertype=abstract.

Seo H-S, Serra R. 2009. Tgfbr2 is required for development of the skull vault. Dev Biol. 334(2):481–90. doi:10.1016/j.ydbio.2009.08.015. http://www.pubmedcentral.nih.gov/articlerender.fcgi?artid=2753698&tool=pmcentrez&rendertype=abstract.

Shukunami C, Yoshimoto Y, Takimoto A, Yamashita H, Hiraki Y. 2016. Molecular characterization and function of tenomodulin, a marker of tendons and ligaments that integrate musculoskeletal components. Jpn Dent Sci Rev. 52(4):84–92. doi:10.1016/j.jdsr.2016.04.003. http://www.ncbi.nlm.nih.gov/pubmed/28408960.

Sijaona A, Luukko K, Kvinnsland IH, Kettunen P. 2012. Expression patterns of Sema3F, PlexinA4, -A3, Neuropilin1 and -2 in the postnatal mouse molar suggest roles in tooth innervation and organogenesis. Acta Odontol Scand. 70(2):140–148. doi:10.3109/00016357.2011.600708. http://www.ncbi.nlm.nih.gov/pubmed/21815834.

Taylor PE, Byers MR, Redd PE. 1988. Sprouting of CGRP nerve fibers in response to dentin injury in rat molars. Brain Res. 461(2):371–6. http://www.ncbi.nlm.nih.gov/pubmed/3263169.

de Vega S, Iwamoto T, Nakamura T, Hozumi K, McKnight DA, Fisher LW, Fukumoto S, Yamada Y. 2007. TM14 is a new member of the fibulin family (fibulin-7) that interacts with extracellular matrix molecules and is active for cell binding. J Biol Chem. 282(42):30878–88. doi:10.1074/jbc.M705847200. http://www.ncbi.nlm.nih.gov/pubmed/17699513.

Wang Y, Cox MK, Coricor G, MacDougall M, Serra R. 2013. Inactivation of Tgfbr2 in Osterix-Cre expressing dental mesenchyme disrupts molar root formation. Dev Biol. 382(1):27–37. doi:10.1016/j.ydbio.2013.08.003. http://www.ncbi.nlm.nih.gov/pubmed/23933490.

Woods SM, Mountjoy E, Muir D, Ross SE, Atan D. 2018. A comparative analysis of rod bipolar cell transcriptomes identifies novel genes implicated in night vision. Sci Rep. 8(1):5506. doi:10.1038/s41598-018-23901-6. http://www.ncbi.nlm.nih.gov/pubmed/29615777.

Zhao H, Oka K, Bringas P, Kaartinen V, Chai Y. 2008. TGF-beta type I receptor Alk5 regulates tooth initiation and mandible patterning in a type II receptor-independent manner. Dev Biol. 320(1):19–29. doi:10.1016/j.ydbio.2008.03.045. http://www.pubmedcentral.nih.gov/articlerender.fcgi?artid=3629921&tool=pmcentrez&rendertype=abstract.

