## supplemental files for "Tgfbr2 in dental pulp cells guides sensory innervation in developing teeth"

### Abbreviations

| | |
|---|---|
| β3T | Beta 3 tubulin |
| BDNF | Brain-derived neurotrophic factor |
| BSA | Bovine serum albumin |
| DP | Dental pulp |
| Dspp | Dentin sialophosphoprotein |
| Fbln7 | Fibrilin 7 |
| Gap43 | Growth associated protein 43 |
| GDNF | Glial cell-derived neurotrophic factor |
| M1 | First (mandibular) molar |
| mTmG | Membrane Tomato membrane GFP |
| Mypn | Myopalladin |
| NGF | Nerve growth factor |
| Osx-Cre | Osterix-Cre-GFP mouse model |
| qPCR | quantitative (real-time) polymerase chain reaction |
| Slitrk6 | SLIT and NTRK Like Family Member 6 |
| TGFβ | Transforming growth factor beta |
| Tgfbr2 | TGFβ receptor 2 |
| $Tgfbr2^{cko}$ | Tgfbr2 conditional knockout |
| TG(N) | Trigeminal (neuron) |
| Tnmd | Tenomodulin |

### Supplemental Table 1 – primer sequences

| Gene | Sequence/Verification method |
|---|---|
| B2 MG F | TTC AGT CGT CAG CAT GGC TC |
| B2MG R | TGA GGG GTT TTC TGG ATA GCA |
| Fbln 7 F | ATC TCC CTT TCA GTG CGA GC |
| Fbln 7 R | GAG CGT GAT GGG TGT CTT CA |
| Gap43 F | GAG GAG CCT AAA CAA GCC GA |
| Gap43 R | TCG TCT ACA GCG TCT TTC TCC |
| Gphn F | TCA TGG CAA ATC ATG GGC AG |
| Gphn R | CAC CTG GAC TGG ACT TTT GGT |
| Mypn F | TCC AGC AGT GTC AGA GTC CT |
| Mypn R | CTC GAC TTT CGG AGA TGG GG |
| Slitrk6 F | TAA GGG CTC AGT GTT TCG CC |
| Slitrk6 R | ATG ATT GGA TCT GAC TCT GTA AAG |
| Tnmd F | TGT ACT GGA TCA ATC CCA CTC T |
| Tnmd R | GCT CAT TCT GGT CAA TCC CCT |
| Tgrbr2e2 F | TTA ACA GTG ATG TCA TGG CCA GCG |
| Tgfbr2e2 R | AGA CTT CAT GCG GCT TCT CAC AGA |
| Wnt10a F | CTTC AGC CGA GGT TTT CGA GAG |
| Wnt10 R | TTC AGT TTA CCC AGA GCG CA |

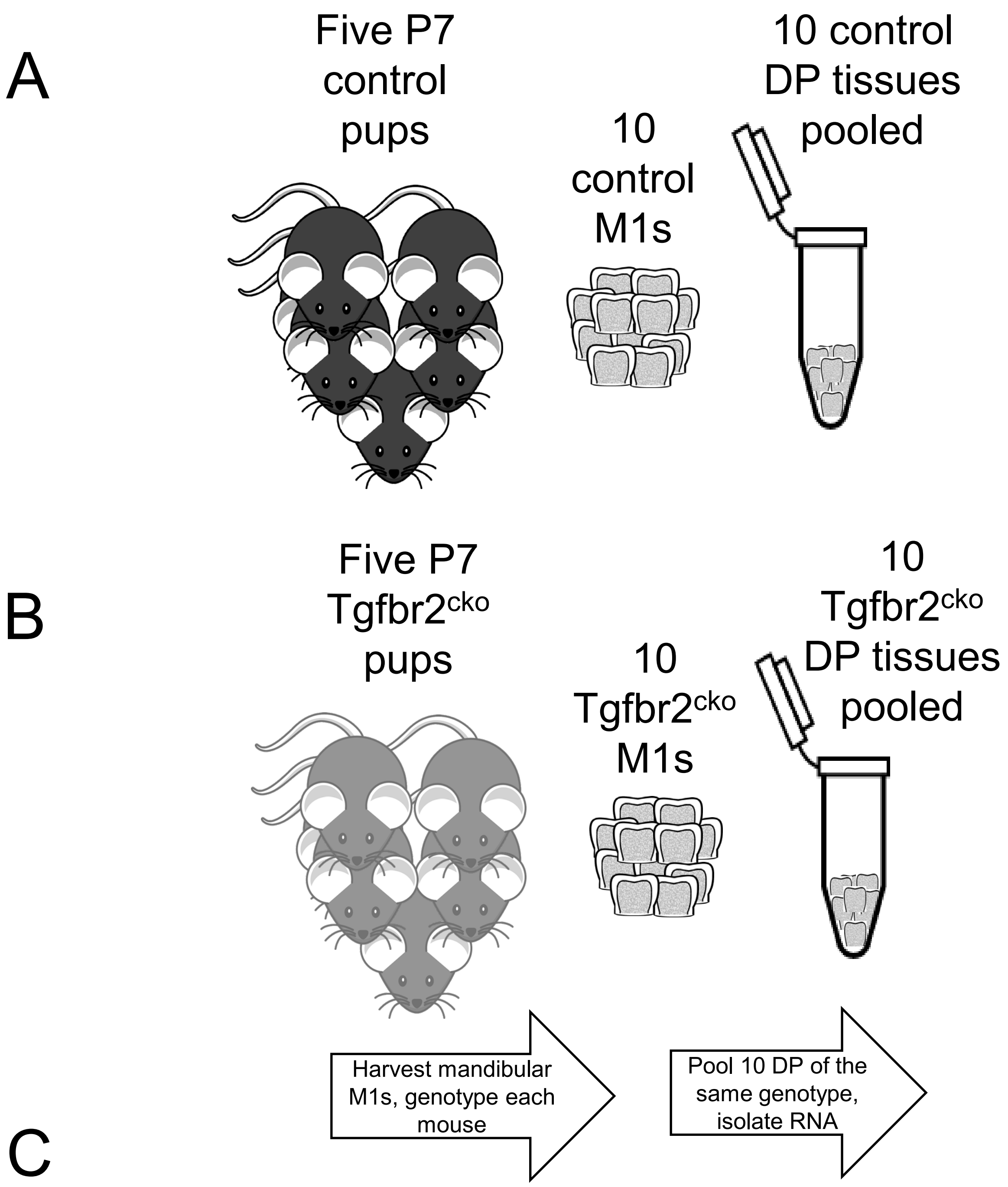


| | n=1 | | | | | n=2 | | | | | n=3 | | | | | n=4 | | | | | n=5 | | | | | n=6 | | | | |
|---|---|---|---|---|---|---|---|---|---|---|---|---|---|---|---|---|---|---|---|---|---|---|---|---|---|---|---|---|---|---|
| Control mice | 1 | 2 | 3 | 4 | 5 | 6 | 7 | 8 | 9 | 10 | 11 | 12 | 13 | 14 | 15 | 16 | 17 | 18 | 19 | 20 | 21 | 22 | 23 | 24 | 25 | 26 | 27 | 28 | 29 | 30 |
| Tgfbr2$^{cko}$ mice | 31 | 32 | 33 | 34 | 35 | 36 | 37 | 38 | 39 | 40 | 41 | 42 | 43 | 44 | 45 | 46 | 47 | 48 | 49 | 50 | 51 | 52 | 53 | 54 | 55 | 56 | 57 | 58 | 59 | 60 |

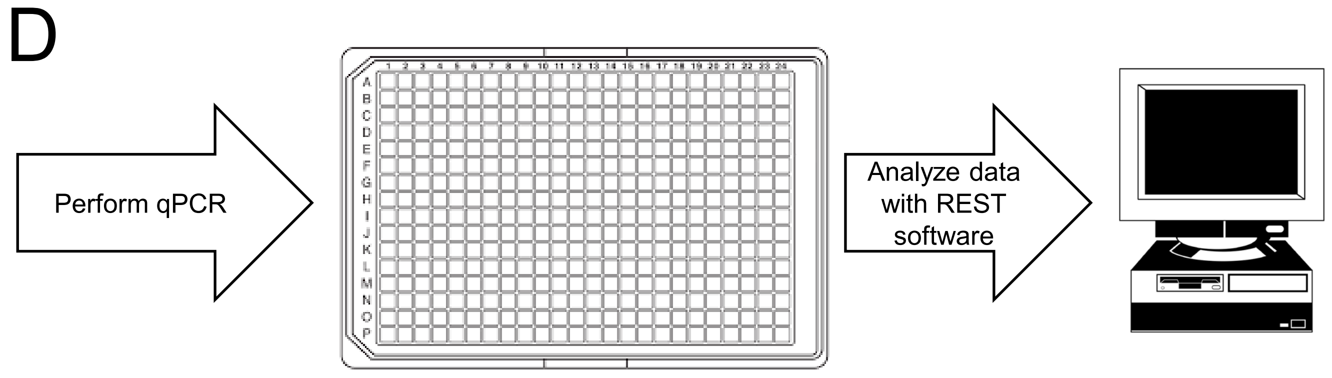


**Supplemental Figure 1.** Schematic of DP RNA collection and processing for qPCR analysis. (A) 5 samples of control are pooled. (B) 5 samples of Tgfbr2$^{cko}$ samples are pooled. (A+B) are considered an n of 1. (C) Chart demonstrating 12 independently pooled samples for 6 separated analyses. Each mouse/sample is assigned a number. (D) qPCR is performed for 6 biological replicates for each genotype.

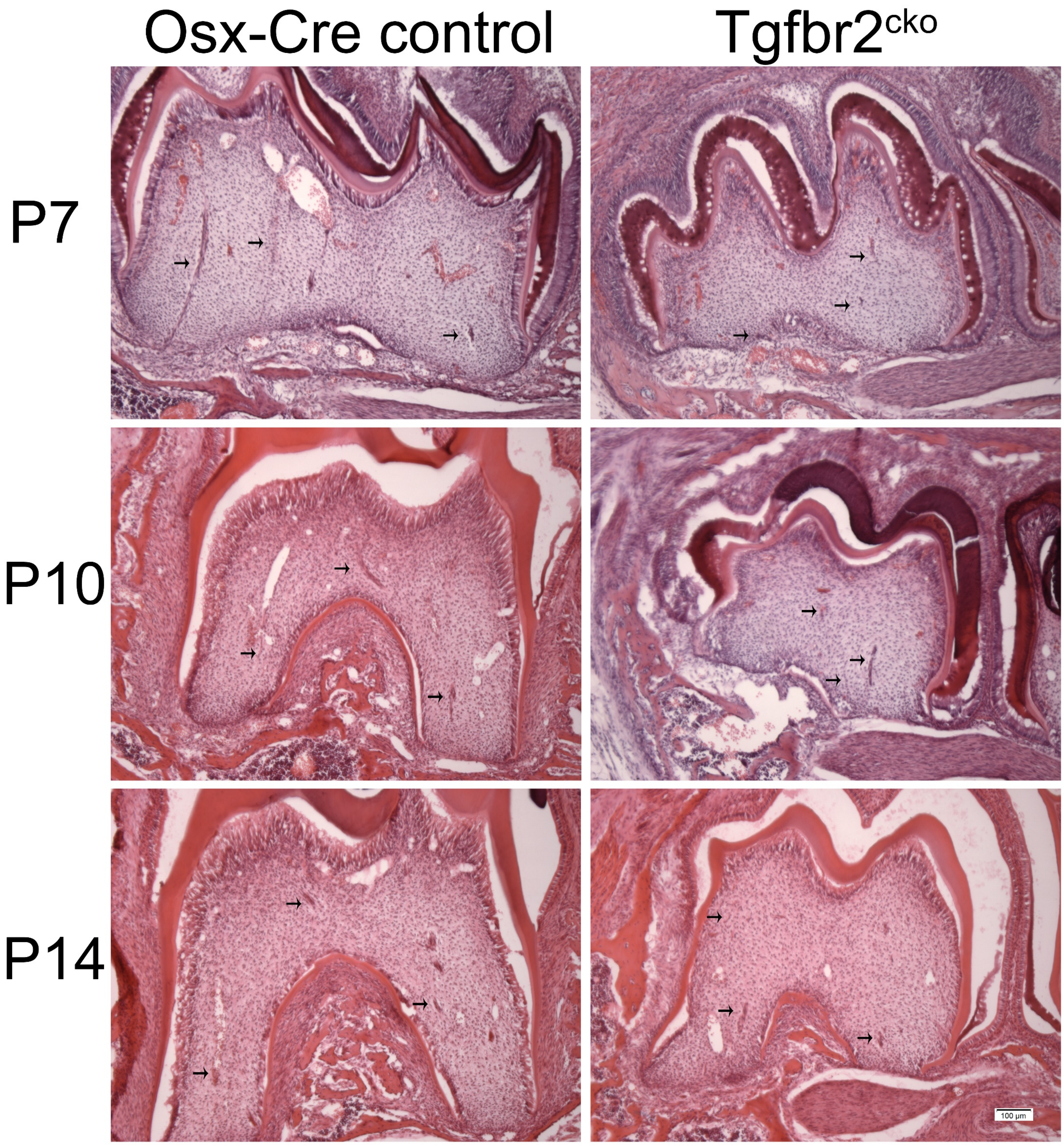


**Supplementary Figure 2:** Nerve structures in postnatal teeth appear to be reduced in Tgfbr2$^{cko}$ mice. Representative images of H&E-stained mandibular M1s from postnatal days 7, 10, and 14. The black arrows in each image indicate hypothesized neuronal structures. Several other structures are present but are not indicated to prevent image crowding. N=3 for each genotype. Scale bar = 100 μm.

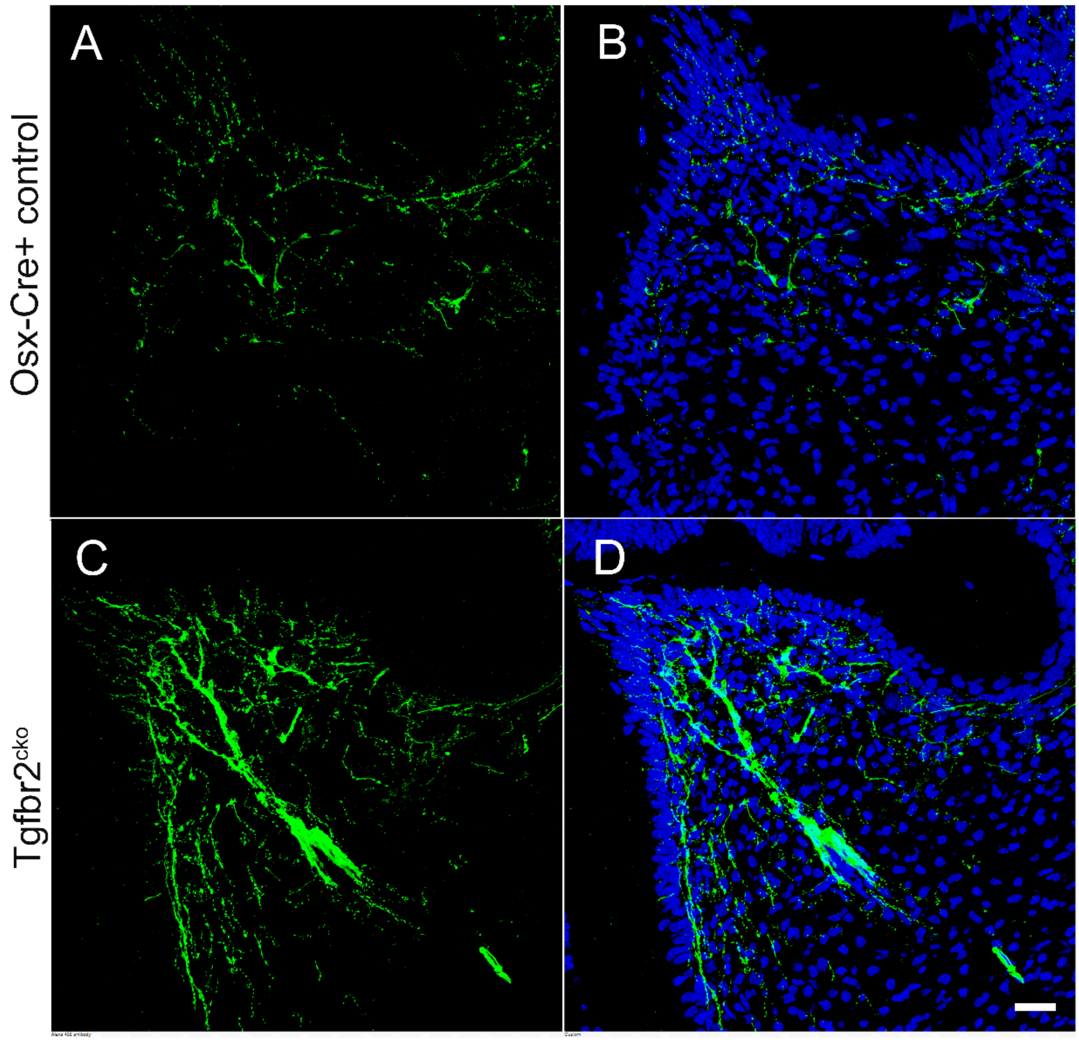


**Supplemental Figure 3.** Gap43 expression is increased in Tgfbr2$^{cko}$ mice. Z-stacks (10 μm thick) were collected and presented in maximum projection mode to examine Gap43 expression. Representative confocal images of control (A, B) and Tgfbr2$^{cko}$ (C, D) dental pulp demonstrated more Gap43 in the mutant mice. The scale bar is 25 μm. n=5 per genotype.
